# Pan-Cancer Biomarker miR-151a Regulates p21 Partially Through p53

**DOI:** 10.1101/2023.01.30.526369

**Authors:** Jessica S. Kurata, Ren-Jang Lin

**Author notes:** **Corresponding authors:** Jessica S. Kurata, Bioinformatics, Guardant Health Inc. Redwood City, CA 94063., Ren-Jang Lin, Beckman Research Institute, 1500 E. Duarte Road, Duarte, CA 91010.

## Abstract

Dysregulation of microRNAs (miRNAs) has been associated with a variety of cancers. We previously identified miR-151a as a potential cell fitness-regulating microRNA using a miRNA-targeted CRISPR-Cas9 genetic screen. In this study, we created mutant cell clones with loss of miR-151a expression and verified miR-151a mutations indeed decreased cell growth. In miR-151a mutant cells, there was an increase in the fraction of cells in the G1-phase of the cell cycle. This increase in G1 cells corresponded to an increase in p53 (TP53) and p21 (CDKN1A) protein levels. Both strands of miR-151a could suppress p53; miR-151a-3p was able to directly suppress p53 expression, but the miR-151a-5p suppression of p53 apparently was indirect. Re-expression of miR-151a-5p in the mutant cells significantly decreased the p53 and p21 protein levels as well as the percentage of cells in G1, while re-expression of miR-151a-3p ironically had a modest effect. These results suggest that both the 5p and 3p strands as well as additional factors are involved in the regulation of p53/p21 and the cell cycle by miR-151a. We also analyzed the TCGA database and discovered that increased miR-151a expression occurs in many tumor types; furthermore, there was an inverse correlation between miR-151a and p21 expression, and high miR-151a expression was often associated with poor overall survival. Taken together, results from this study identify a previously underappreciated role of miR-151a in cancer through regulation of the cell cycle, and they also suggest inhibiting the less abundant 5p may be more important than inhibiting the more abundant 3p of miR-151a for therapeutic considerations.

## INTRODUCTION

MicroRNAs (miRNAs) are short, single-stranded RNA molecules that post-transcriptionally regulate the expression of target mRNAs. Mature microRNAs are produced from stem-loop-containing primary transcripts through a series of cleavage steps. The stem-loop can be processed to produce both 5p and 3p mature miRNAs, though some primary miRNAs show strong bias to a single mature miRNA. The two mature miRNAs from a single primary transcript may have similar roles, such as miR-223-5p and -3p that both inhibit bladder cancer migration ^1^, or they may have opposite roles, such as the inhibition of cellular proliferation by miR-28-5p and the promotion of proliferation by miR-28-3p in colorectal cancer ^2^.

We previously identified 44 putative pro-fitness microRNAs in HeLa cells using a genome-wide CRISPR-Cas9 screen ^3^. Cell fitness in the screen infers the ability of the cell to compete in cell growth; thus, inactivation of a pro-fitness microRNA by CRISPR shall lead to reduced cell proliferation. We decided to further investigate miR-151a, as it has been shown to be overexpressed in cervical tumor samples but its role in cancer is not well understood. miR-151a is produced from intron 22 in the focal adhesion kinase gene (*FAK*/*PTK2*), and there is a strong correlation between miR-151a and FAK expression seen in tissue and cell line samples ^4^. The miR-151a primary transcript produces two mature miRNAs, with miR-151a-3p often expressed at a higher level than miR-151a-5p.

That miR-151a-5p overexpression increases migration and invasion has been shown in hepatocellular carcinoma ^4^, prostate cancer ^5^, non-small cell lung cancer (NSCLC) ^6^, and gastric cancer cells ^7^. miR-151a-5p overexpression increases cell growth and colony formation in gastric cancer ^7^ and NSCLC ^6^ cells, but it does not change cellular proliferation in hepatocellular carcinoma ^4^ or prostate cancer ^5^ cells. Several mRNAs have been reported to be targets of miR-151a-5p, such as RhoGDIA (encoded by the *ARHGDIA* gene) ^4, 5^, CASZ1 ^5^, IL1RAPL1 ^5^, SOX17 ^5^, N4BP1 ^5^, and TP53 ^7^. Overexpression of miR-151a-3p in gastric cancer cells is also found to increase migration and invasion ^7^. miR-151a-3p is known to regulate CDH1 ^6^, which encodes E-cadherin. Previous work has suggested a role of miR-151a-5p and 3p in the migration and invasion of cancer cells; however, ability of miR-151a to regulate cellular fitness is less well defined.

To further determine the role of miR-151a in cellular fitness or cell growth, we used CRISPR-Cas9 to generate deletions in the miR-151a gene in HeLa cells and isolated mutant colonies. We verified miR-151a mutations indeed resulted in decreased cellular fitness and found it also increased the fraction of cells in the G1-phase of the cell cycle. When the mRNA expression profile of the miR-151a mutant cell lines was examined, we found up-regulation of cell cycle, p53, and TGF-β pathway genes. We further investigated the link between miR-151a and p53 and found miR-151a-3p, but not 5p, can directly target the p53 3’UTR in a reporter assay. We also found miR-151a mutation increased the protein level of p53 as well as the level of p21, a downstream transcriptional target of p53. When miR-151a-3p or -5p was strand-specifically re-expressed in miR-151a mutant cells, miR-151a-3p overexpression unexpectedly only resulted in a modest decrease in p53 and p21 protein levels. miR-151a-5p overexpression, on the other hand, resulted in a significant decrease in p53 and p21 protein levels and a decrease in the percentage of cells in the G1 phase of the cell cycle. Thus, it appears that although miR-151a-3p directly suppressed p53/p21, 5p exhibited stronger suppression perhaps through targeting other factors that resulted in p53/p21 down-regulation. We also investigated the clinical relevance of this regulation by examining the expression of miR-151a across patient tumor samples in the TCGA database. In cases where tumor and paired normal samples were available, miR-151a was found to be up-regulated in thirteen out of fourteen cancers. We found a significant negative correlation between miR-151a expression and the p21 mRNA level in approximately half of the thirty-two cancers examined, and only in one case that the opposite correlation was observed. Moreover, we discovered an association between miR-151a overexpression and poor overall survival in seven cancers. Taken together, our results indicate that an elevation of miR-151a expression can lead to p53/p21 suppression and cell proliferation, and that miR-151a may be an underappreciated pan-cancer biomarker.

## MATERIALS AND METHODS

### Cell Culture & HeLa-Cas9/HeLa-Cas9 GFP^+^ Creation

HeLa cells were grown in DMEM with 10% FBS at 37°C and 5% CO_2_. The HeLa-Cas9 cell line was isolated as previously described [citation]. To create the HeLa-Cas9 GFP^+^ cell line, HeLa-Cas9 cells were transduced with HIV_7_/CMV-GFP lentivirus. Cells with high GFP expression were sorted out using an Aria SORP (BD Biosciences) to produce the HeLa-Cas9 GFP^+^ cell line.

### Vectors

pLX-sgRNA (Addgene plasmid # 50662) and pCW-Cas9 (Addgene plasmid # 50661) were a gift from Eric Lander & David Sabatini^8^. HIV_7_/CMV-GFP and HIV_7_/PGK-neo were a gift from Tammy Chang and Jiing-Kuan Yee. U6-NX and psiCHECK-2.2 were a gift from Guihua Sun^9^. We modified pLX-sgRNA to allow sgRNA insertion by ligation as previously described^3^. The 3’-UTR of TP53 and TGFB3 were PCR amplified from HeLa genomic DNA using the TP53-UTR and TGFB3-UTR primers (Supplemental Table S1). The PCR product was gel purified using the Nucleospin Gel and PCR Clean-Up Kit (TaKaRa Bio Inc.) and cloned in between the XhoI and SalI sites of psiCHECK-2.2 using InFusion cloning (TaKaRa Bio Inc.). To create mutations to the predicted miR-151a binding sites, the QuikChange Lightning (Agilent) was used. The sequences for the TP53-3p-SDM, TP53-5p-SDM, TP53-5p2-SDM, and TGFB3-5p-SDM primers are given in Supplemental Table S1.

### miR-151a Mutant Cell Line Generation

miR-151a targeting sgRNA oligos (sg-151a; Supplemental Table S1) were annealed and then ligated between the BfuAI sites of the pLX-sgRNA-BfuAI-2k vector. Lentivirus was produced and HeLa-Cas9 cells were transduced. After selecting for transduced cells using 5 µg/mL blasticidin, Cas9 expression was induced using 2 µL doxycycline for one week. The cells were then plated on 96-well plates diluted to 1 cell per well. Single cell colonies were screened for miR-151a deletion using fragment analysis using the previously described protocol^10^. Briefly, PCR fragments were FAM labeled using the miR-151a-FA primers and a M13(−21)-FAM primer (Supplemental Table S1). The size of these fragments was determined by capillary electrophoresis. Those colonies with no peak at the wild type length were then Sanger sequenced to determine the exact deletions present.

### GFP Competitive Growth Assay

To determine the relative growth rate of HeLa-Cas9 GFP^+^ cells and another cell line, 500,000 cells from each cell line were mixed. Half of these cells were used to measure the day 0 ratio of GFP^+^ to GFP^-^ cells and the other half were plated. Every three days for 12 days the cells were split and the percentage of GFP^-^ cells was determined using flow cytometry (CyAn ADP, Beckman Coulter). The percentage of GFP^+^ cells was determined using FlowJo v10 (FlowJo, LLC). The % of GFP^-^ cells was normalized using:

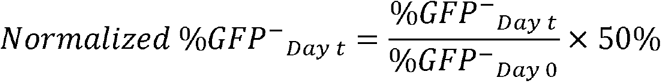

### mRNA-sequencing

mRNA was extracted from ∼5×10^6^ cells using the miRNeasy Mini Kit (QIAGEN). Library preparation and sequencing was conducted by City of Hope Integrative Genomics Core. Library construction of 300 ng total RNA for each sample was made using KAPA Stranded mRNA-Seq Kit (Illumina) (Kapa Biosystems), using 10 cycles of PCR amplification. Libraries were purified using AxyPrep Mag PCR Clean-up kit (Thermo Fisher Scientific). Each library was quantified using a Qubit fluorometer (Life Technologies) and the size distribution assessed using the 2100 Bioanalyzer (Agilent Technologies).

Sequencing was performed on an Illumina Hiseq 2500 (Illumina) instrument using the TruSeq SR Cluster Kit V4-cBot-HS (Illumina) to generate 51 bp single-end reads sequencing with v4 chemistry. Quality control of RNA-Seq reads was performed using FastQC. The raw reads were then aligned using the splicing aware aligner TopHat (version 2.0.4), which relies on Bowtie2 (version 2.0.0-beta5)^11^ for read alignment. The gene locations were downloaded from Ensembl (assembly 38, release 88). Since the Bowtie2 index was built using hg38, the chromosome names in the gtf file were converted from Ensembl to UCSC naming conventions. Once the reads had been aligned, they were counted using the HTSeq (version 0.7.2)^12^ function HTSeq-count. Differential expression analysis was then performed using the R (version 3.3) package EdgeR (version 3.16.5)^13, 14^.

### miR-151a Strand-specific Overexpression

To overexpress miR-151a in a strand specific manor, sli-miRNA constructs^9^ were created by annealing the sli-miR1515p, sli-miR1513p, or sli-Ctrl (Supplemental Table S1) oligos. The annealed oligos were then ligated between the BglII and XhoI sites of U6-NX.

To create lentivirus, the U6 promoter and sli-miRNA region of the vectors were amplified by PCR using the sli-InFus (Supplemental Table S1) primers, gel purified using the Nucleospin Gel and PCR Clean-Up Kit (TaKaRa Bio Inc., Kusatsu, Japan), and then cloned into the XbaI site of pHIV_7_/PGK-neo using InFusion cloning (TaKaRa Bio Inc.) to create the pHIV_7_/sli-miR-151a-3p-PGK-neo, pHIV_7_/sli-miR-151a-5p -PGK-neo, and pHIV_7_/sli-Ctrl-PGK-neo constructs which were used to create stable re-expression cell lines.

### TCGA Analysis

Level 3 data for all 33 TCGA projects was downloaded using the GDC tool. GBM was excluded from analysis as no tumor samples have miRNA-sequencing data. In all cases, the miRNA expression in primary tumor samples was examined and metastatic samples were disregarded. For tumor versus normal analysis, projects with more than 10 paired tumor and normal samples were selected. The raw read counts for those samples with paired data were collected and analyzed using DESeq2^15^.

To determine the association between miRNA expression and patient survival, the 25% of patients with the highest or lowest miR-151a-5p or -3p expression were identified using the read per million normalized read counts. The overall survival of patients in the highest 25% was compared to the patients in the lowest 25% using the log-rank test.

The correlation between read per million normalized read counts for miR-151a and trimmed mean of M values (TMM) normalized^16^ read counts for CDKN1A was determined using Pearson’s correlation.

Additional information is available in the Supplemental Materials and Methods.

## RESULTS

### Isolation of miR-151a Mutant Cell Lines

To investigate the function of miR-151a, we constructed miR-151a mutant HeLa cells using CRISPR-Cas9. Five miR-151a-targeting sgRNAs have been included in the competitive cell growth screen in our previous genome-wide cell fitness study (Fig. 1A) ^3^. We chose sgRNA sg22705 as the mutagen, which would induce Cas9-cleavage at 18 bp downstream of the Drosha cleavage site, in the CNNC motif necessary for SRp20/p72 binding and efficient Drosha cleavage (Fig. 1B) ^17, 18^. Because sg22705 induced cleavage near but not at the mature miR-151a sequence, it might generate partial and complete loss-of-function alleles that would result in knockdown and knockout mutants.

**Figure 1.**
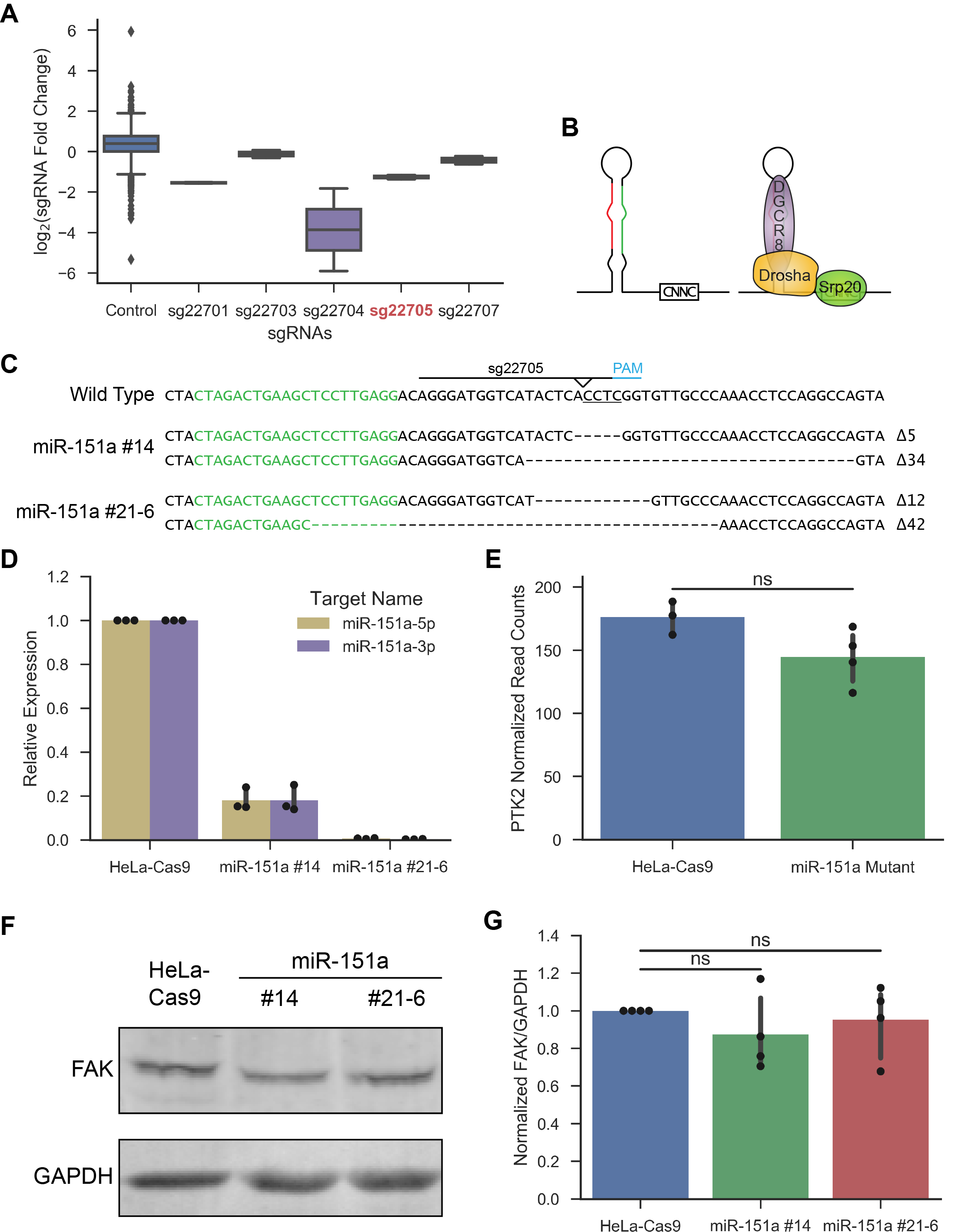
Isolation of miR-151a mutant colonies. **A**, Box plot showing a negative fold change of the miR-151a-targeting sgRNAs versus the control sgRNAs in two replicates of the LX-miR screen in HeLa cells. The box shows the quartiles, the lines show the range and dots represent outliers. SgRNA 22705 labeled in red was used for knockout experiments. **B**, Diagram of pre-miRNA with the CNNC sequence and the binding of Drosha, DGCR8, and Srp20 to the pre-miRNA. The 5p strand is shown in red, while the 3p strand in green. **C**, DNA sequence of the miR-151a region of wild type or miR-151a mutant colonies. The green sequence is miR-151a-3p and the line indicates the sg22705 target sequence including the Cas9 cleavage site. The CNNC motif is underlined. Dotted lines indicate deleted nucleotides. **D**, qRT-PCR of miR-151a-3p and - 5p expression in parental and miR-151a mutant colonies. Data points show three independent experiments. Error bars represent 95% confidence interval CI. **E**, PTK2/FAK expression of parental and mutant colonies was estimated from RNA-Seq data. Poly(A)RNA was collected twice from each miR-151a mutant colony and three times from HeLa-Cas9 parental cells. Statistics were calculated using EdgeR (ns: FDR > 0.05). Data points are TMM normalized read counts. **F**, Representative western blot of FAK protein levels in HeLa-Cas9 parental and miR-151a mutant cells. GAPDH was used as a loading control. **G**, Western quantification of FAK protein levels in parental and miR-151a mutant cells. Data points show values from four independent experiments. Error bars represent 95% CI. Statistics calculated using two-tailed paired *t*-test (ns: *p* > 0.05).

HeLa-Cas9 cells, which contained Cas9 under the control of doxycycline-inducible promoter, were transduced with the miR-151a-targeting sg22705 sgRNA. After selecting for transductants, we induced Cas9 expression and isolated single colonies. These colonies were then screened for miR-151a indel mutations using fragment analysis. One homozygous mutant colony, colony #14, was obtained. A heterozygous mutant, colony #21, was further mutagenized in another round of Cas9 induction and colony screening. The second round of colony screening led to the isolation of another homozygous mutant colony #21-6. Sequencing of the miR-151a locus showed that colony #14 had a 5-bp and a 34-bp deletions, and colony #21-6 had a 12-bp and a 42-bp deletions (Fig. 1C). The effect of these mutations on miR-151a-5p and -3p expression was determined by qRT-PCR; miR-151a-3p and -5p expression was almost completely abolished (0.3-0.7% of parental) in mutant miR-151a #21-6, while mutant miR-151a #14 retains approximately 20% expression of both strands (Fig. 1D). We also carried out small RNA sequencing and found a ∼4-fold decrease in miR-151a-3p and -5p expression in mutant miR-151a #14 (Supplemental Fig. S1).

The miR-151a sequence is in imbedded in intron 22 of the FAK/PTK2 gene, which is 2.5 kilobase pairs from the upstream exon and 15 kbp from the downstream exon. It was unlikely that the small indel mutations would directly affect the expression of PTK2, but nonetheless it needed to be analyzed. We carried out RNA sequencing and found there was a slight decrease in FAK/PTK2 RNA reads in the miR-151a mutant cells, but the difference was not significant (Fig. 1E). We also measured the protein levels of FAK/PTK2 by immunoblotting and again found a slight, not statistically significant, decrease in the mutant cells (Fig. 1F and 1G). Thus, it should be rather safe to assume that the molecular and cellular consequences we would be measuring were most likely linked to the loss of miR-151a and not to the subtle changes of FAK/PTK2.

### miR-151a Mutation Decreased Fitness and Increased G1-phase Cell Percentage

We investigated the impact of miR-151a mutations on cell proliferation and fitness using a competitive growth assay. We marked the parental HeLa-Cas9 cells with a GFP gene introduced by a lentiviral vector and used them in the competition assay. The miR-151a mutant cells, which were GFP^-^, were mixed with the aforementioned wild-type miR-151a GFP^+^ cells, and the mixed cell population was cultured; the relative fraction of GFP^-^ cells in the population over time was determined by flow cytometry (Fig. 2A). The percentage of GFP^-^ miR-151a mutant cells drastically decreased over time, indicating the miR-151a mutant cells had decreased fitness relative to the miR-151a wild-type cells (Fig. 2B). The control experiment was to co-culture the parental HeLa-Cas9 GFP^-^ cells with HeLa-Cas9 GFP^+^ cells, which showed a slight increase of the GFP^-^ parental cells over time. Thus, it appeared that either the expression of GFP had some negative effect on cell fitness, or the expression of GFP in some GFP+ cells was silenced over time. Nevertheless, this growth competition experiment confirmed the importance of miR-151a for maintenance of cellular fitness, and was consistent with what was seen in our genome-wide screen, where cells containing miR-151a-targeting sgRNAs gradually disappeared from the pool ^3^.

**Figure 2.**
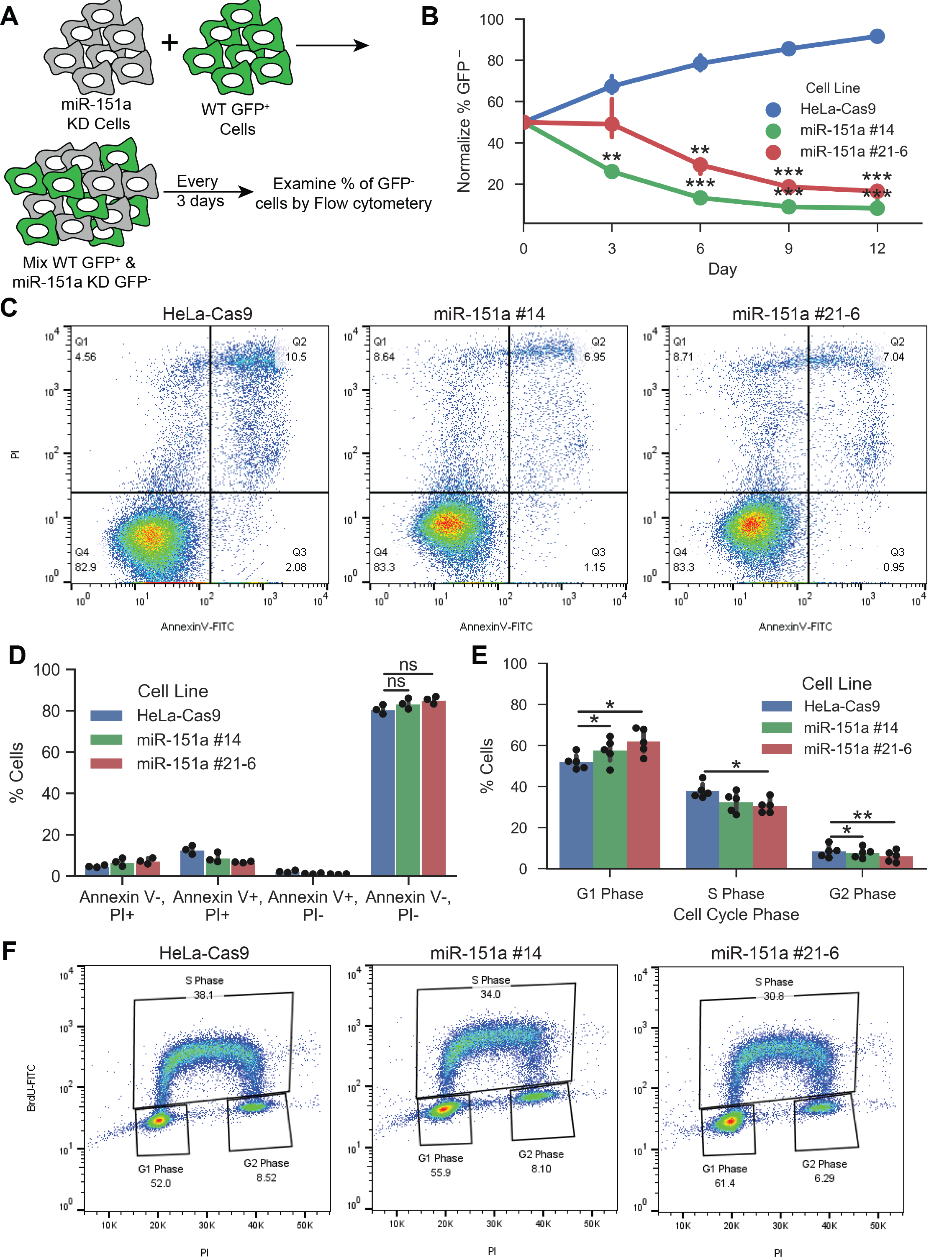
miR-151a mutant colony phenotypes. **A**, Diagram of competitive growth assay between GFP^-^ testing cells versus GFP^+^ control cells. **B**, The percentage of GFP^-^ cells over time was determined using flow cytometry. Points represent the mean % GFP^-^ cells for three independent experiments. **C**, Flow cytometry apoptotic assays. HeLa-Cas9 parental and miR-151a mutant cells were analyzed by Annexin V and propidium iodide (PI) staining. **D**, Quantification of Annexin V and PI staining of HeLa-Cas9 and miR-151a mutant cells was determined. Data points represent three independent replicates. **E**, Cell cycle analysis. HeLa-Cas9 and miR-151a mutant cells were incubated in media containing BrdU, then co-stained with anti-BrdU and PI and analyzed by flow cytometry to determine the cell cycle distribution. Data points represent five independent experiments. **F**, Representative image of anti-BrdU and PI flow cytometry analysis. For **B, D**, and **E**, statistics were calculated using two-tailed paired *t*-test (ns: *p* > 0.05; * *p* < 0.05; ** *p* < 0.01; *** *p* < 0.001). Error bars represent 95% confidence interval.

We examined the impact of miR-151a mutation on apoptosis, and found no significant difference in the percentage of viable cells between HeLa-Cas9 parental cells and miR-151a mutant cells (Fig. 2C and 2D). We then examined the cell cycle in these cells, and found miR-151a mutant cells showed a significant increase in percentage of G1 phase cells and decreases in percentage of G2 and S phase cells (Fig. 2E and 2F). We also examined the impact of miR-151a mutation on the ability of HeLa cells to form colonies and found the number of colonies formed by the miR-151a mutants and the number by the wild-type were not significantly different (data not shown). These phenotype analyses indicated that miR-151a mutation did not affect the viability or colony formation ability of HeLa cells, but it did significantly decrease the fitness of the cell and increase the portion of cells in the G1 phase of the cell cycle.

### miR-151a Mutation Resulted in Up-regulation of genes in the TGF-β, p53, and Cell Cycle Pathways

To determine the molecular mechanisms by which miR-151a regulates these phenotypes, we examined the mRNA expression profile in parental and mutant cells by RNA sequencing. By comparing genome-wide mRNA abundance, we identified 1,781 significantly differentially expressed genes (FDR < 0.05) with 740 genes up-regulated and 1,041 genes down-regulated in the miR-151a mutants (Supplemental Fig. S2A). Pathway analysis of these genes using either Gene Set Enrichment Analysis ^19, 20^ or Ingenuity Pathway Analysis ^21^ showed an enrichment in cell cycle, p53, and TGF-β related pathways among those genes up-regulated after miR-151a mutation (Supplemental Fig. S2B, S2C, and S2D). One the other hand, the pathways enriched among down-regulated genes had little overlap between different analysis methods. Since microRNA is mostly functioning by suppressing gene expression, we therefore focused on the pathways linked to the up-regulated (derepressed) genes associated with the loss-of-function mutation. The enrichment of cell cycle-related genes agreed with the change in cell cycle seen in miR-151a mutant cells. As the p53 and TGF-β signaling pathways are both known to regulate cellular fitness and the cell cycle, these pathways were further investigated.

### Predicted miR-151a Targets

To determine through which genes miR-151a regulates the p53, cell cycle, and TGF-β pathways, we searched for predicted miR-151a-5p or -3p targets using TarBase ^22^, microT-CDS ^23, 24^, miRanda-mirSVR ^25^ and TargetScan ^26^. We then analyzed our mRNA-seq data to identify which target genes predicted by at least one method had significantly increased expression in the miR-151a mutants, and 121 miR-151a-3p targets and 64 miR-151a-5p targets were identified as such. Among those genes, 17 miR-151a-3p and 9 miR-151a-5p predicted targets have Gene Ontology GO terms related to the p53, TGF-β, or cell cycle pathways (Supplemental Table S2). Of these, the predicted miR-151a-3p target TP53 was of special interest, because p53 is a known regulator of the cell cycle and increased p53 expression can result in G1 arrest ^27^. TP53 has previously been shown to be a miR-151a target in gastric cancer cells ^7^. HeLa cells are HPV18 positive and express the HPV-E6 protein ^28^, which targets the p53 protein for ubiquitination and degradation ^29^. This E6-induced degradation effectively inhibits the activity of p53 in HeLa cells and likely also in cervical cancer patients, as 95% of which are HPV positive ^30^. Nonetheless, it is possible that, besides the HPV-E6-mediated proteosomal degradation of p53, miR-151a may play an unappreciated role in suppressing the p53 pathway in miR-151a overexpressing cancer (see below).

### p53 was Verified to be a Direct miR-151a-3p Target

Our mRNA-sequencing results showed a significant increase in the expression of *TP53* in miR-151a mutant cells (Fig. 3A). By immunoblotting, we found p53 protein levels also significantly increased in miR-151a mutant cells (Fig. 3B and 3C). To examine whether p53 is a direct target of miR-151a, we cloned the 3’-UTR of TP53, which contains one predicted miR-151a-3p site and two possibly 5p sites, into a dual luciferase reporter vector. For control, we also mutated 3 bases in the seed sequence of the three potential miR-151a binding sites predicted by TargetScan ^26^, RNA22 ^31^, or RNAHybrid ^32^ (Fig. 3D). We then created strand specific miR-151a-5p and -3p expression constructs using a slicer miRNA strategy based on the AGO2-dependent but Dicer-independent processing of miR-451^9^ (Fig. 3E). Co-transfection with the miR-151a-3p, but not -5p expression construct was found to reduce the p53-3’UTR-luciferase activity (Fig. 3F). This suppression was abolished when the predicted miR-151a-3p site was mutated but the suppression was unaffected by mutating either of the two predicted miR-151a-5p binding sites (Fig. 3F). This result suggested miR-151a-3p is able to directly regulate TP53 by binding to its 3’-UTR, while miR-151a-5p appears not to directly regulate TP53.

**Figure 3.**
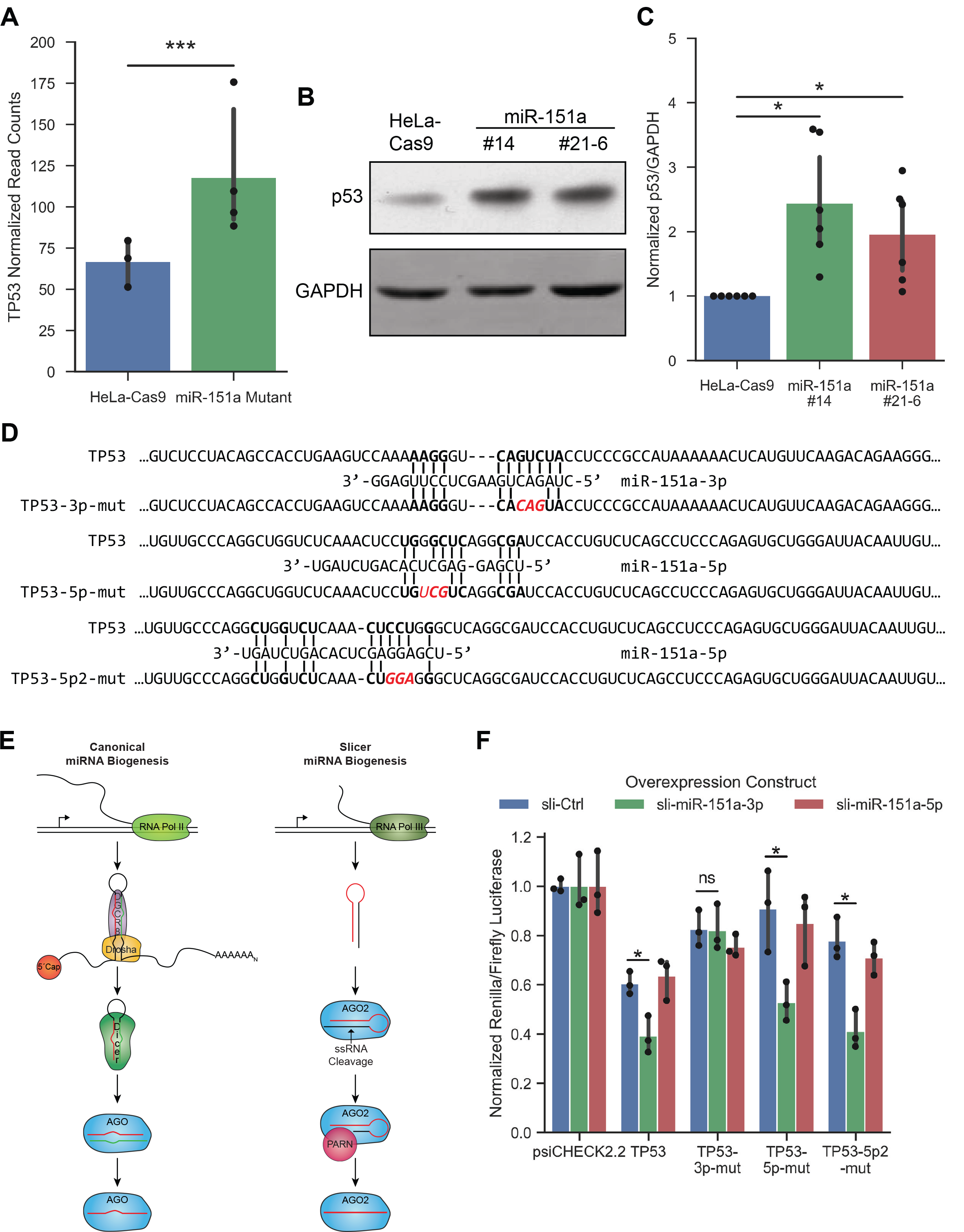
p53 as a direct target of miR-151a-3p. **A**, TP53 expression of parental and mutant colonies was determined by RNA-sequencing. Poly(A)RNA was collected from each cell line and sequencing reads were compared using EdgeR (*** *p* < 0.001). Data points represent TMM normalized read counts. **B**, Representative western blot of p53 levels in HeLa-Cas9 parental and miR-151a mutant cells. GAPDH was used as a loading control. **C**, Quantification of western blot. Data points show values from 6 independent experiments. **D**, Diagram depicting the putative binding sites of miR-151a-5p and -3p in the TP53 3’-UTR as predicted by TargetScan, RNAHybrid, or RNA22. Mutations that were made to interrupt the pairing between the seed sequence of miR-151a and the TP53 3’-UTR are shown in red. **E**, Diagram of canonical and slicer miRNA biogenesis. The mature miRNA sequence is shown in red or in green. **F**, The Renilla to firefly luciferase expression ratio was measured in cells co-transfected with a dual luciferase expressing vector carrying a 3’-UTR variant and a slicer construct expressing miR-151a-5p or -3p. The sli-Ctrl that did not express either miR was used as control. This ratio was normalized to the average value using the control psiCHECK luciferase vector. Data points indicate three independent replicates. For **C** and **F**, statistics were calculated using two-tailed paired *t*-test (ns: *p* > 0.05; * *p* < 0.05). Error bars represent 95% confidence interval.

### Expression of p21 Increased in miR-151a Mutants in a p53 Dependent Manner

To determine the downstream effects the increase in p53 might have on the miR-151 mutant cells, we examined our mRNA-seq data for the expression of known p53-regulated genes. Of the 100 consensus p53 targets identified by ^33^, 16 genes significantly increased in expression after miR-151a mutation (Supplemental Table S3). These 16 include genes involved in the control of DNA damage response, proliferation, apoptosis, and cell cycle regulation. The most significantly up-regulated gene was *CDKN1A*, which encodes p21 (Fig. 4A). p21 is one of the most well studied p53 targets and is known to mediate p53-dependent G1-arrest of the cell cycle ^34-36^. Since we have observed miR-151a mutations resulted in an increased fraction of cells in the G1-phase of the cell cycle (Fig. 2), we decided to investigate the p21 protein levels in miR-151a mutant cells. By western blot, the p21 protein expression levels were found to increased approximately 7 fold after miR-151a mutation (Fig. 4B and 4C).

**Figure 4.**
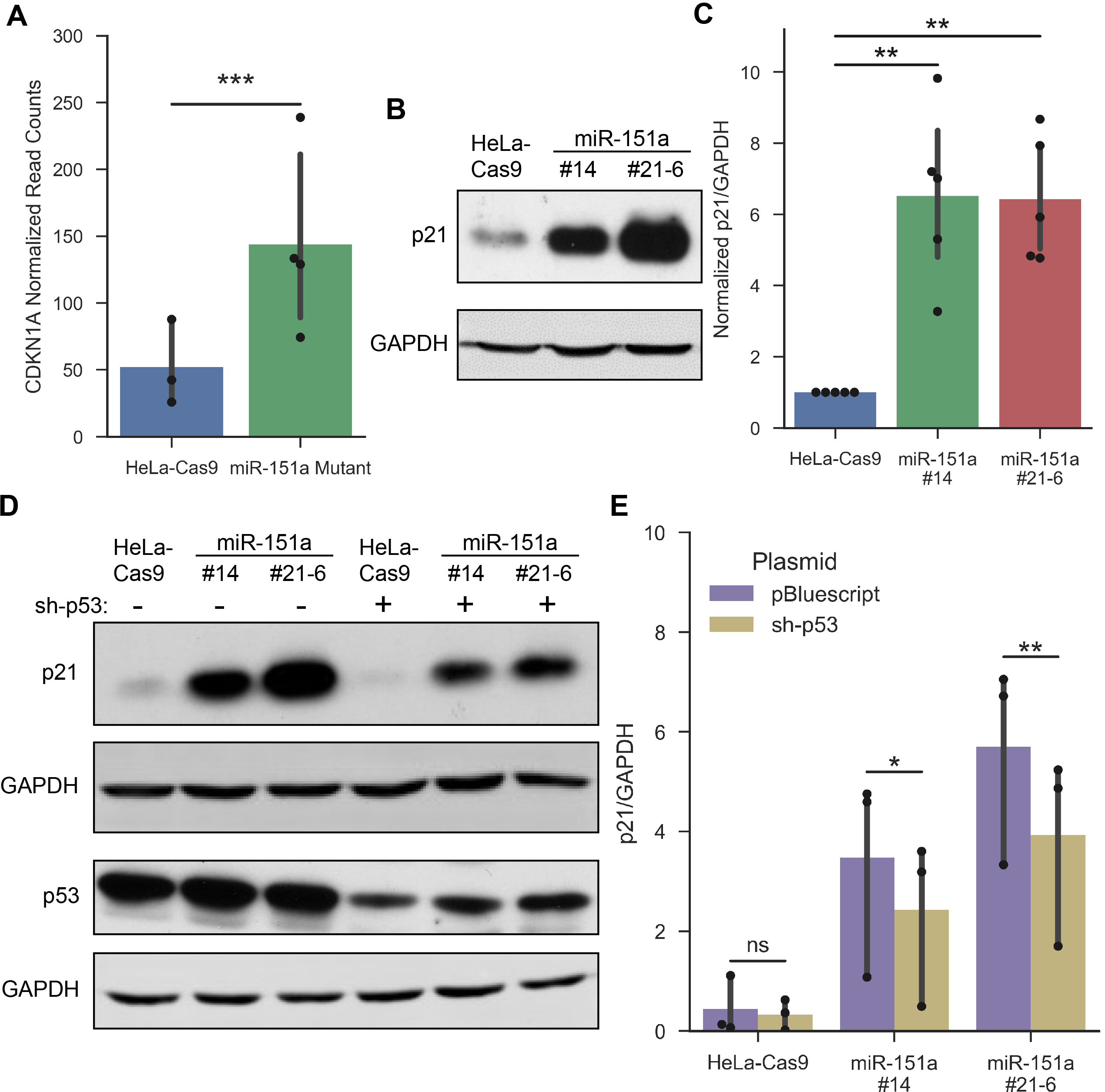
Expression of p21 in cells with miR-151a mutations. **A**, CDKN1A/p21 expression in HeLa-Cas9 parental and miR-151a mutant colonies was deduced from RNA-sequencing normalized read counts. Statistics were calculated using EdgeR (*** *p* < 0.001). **B**, Representative western blot of p21 levels in HeLa-Cas9 parental and miR-151a mutant cell lines. **C**, Quantification of p21 over GAPDH in miR-151a mutants in western blots and normalized to the ratio in HeLa-Cas9. Data points represent five independent experiments. **D**, Representative western blot of parental and miR-151a mutant cells transfected with a shRNA targeting p53 (labeled with +) or empty vector (labeled with -). **E**, Quantification of the ratio between p21 and GAPDH after cells were transfected with sh-p53 or empty vector pBluescript (n=3). For **C** and **E**, statistics were calculated using two-tailed paired *t*-test (ns: *p* > 0.05; * *p* < 0.05; ** *p* < 0.01). Error bars represent 95% confidence interval.

To verify the increase of p21 in miR-151a mutants was p53-dependent, we knocked down p53 by shRNA and examined the p21 expression. Transfection with a p53-targeting shRNA resulted in about 75%-50% decrease in p53 protein levels (Fig. 4D; Supplemental Fig. S3). When p21 protein levels were examined, we found there was a significant decrease in p21 levels upon p53 knockdown in the miR-151a mutant cells (Fig. 4D and 4E). The p21 level in miR-151a mutant cells did not decrease to parental levels after p53 knockdown. This could be due to the incomplete knockdown of p53; however, the p53 levels in miR-151a mutant colonies after transfection with the p53 shRNA was below parental levels (Fig. 4D; Supplemental Fig. S3). It appears that the increase in p21 expression seen after miR-151a mutation may not be entirely p53-dependent.

### Re-expression of miR-151a-5p Reversed Molecular and Cellular Phenotypes

To verify the phenotypical changes observed were due to the mutation of miR-151a and to determine which strand of miR-151a was critical, we transduced the HeLa-Cas9 parental and miR-151a mutant cells with our strand-specific slicer miRNA constructs as described above. The slicer miRNA constructs produced miR-151a-3p, -5p, or a scramble control sequence using an AGO2-dependent biogenesis pathway (Fig. 3E). The expression of miRNAs from these constructs were quite effective; transduction with the sli-miR-151a-3p construct resulted in 3-6 times wild type miR-151a-3p (Supplemental Fig. S4A), while sli-miR-151a-5p transduction resulted in miR-151a-5p expression levels 20-60 fold greater than wild type (Supplemental Fig. S4B). The difference in the fold change of expression after slicer transduction was likely due to the lower basal level of miR-151a-5p expression relative to miR-151a-3p expression. No change in expression of the alternate strand was detected as expected.

We then examined the p53 and p21 protein levels in the miR-151a-3p or -5p re-expressing cell lines. Interestingly, we found re-expression of miR-151a-5p significantly decreased the p53 protein level in transduced miR-151a #21-6 cells (Fig. 5A and 5B). Re-expression of miR-151a-3p also decreased the p53 protein level in transduced miR-151a #21-6 cells; however, this decrease appeared to be more subtle. Since our reporter assay did not support a direct suppression of p53 by miR-151a-5p, this result suggested miR-151a-5p could effectively downregulate p53 levels through an indirect mechanism. In slicer construct-transduced miR-151a #14 cells, the changes in p53 expression were marginal (Supplemental Fig. S5A and S5B); nonetheless, a significant decrease in p21 expression was observed after miR-151a-5p re-expression (Supplemental Fig. S5A and S5C). Thus, a re-expression of miR-151a-5p drastically decreased the amount of p21 protein.

**Figure 5.**
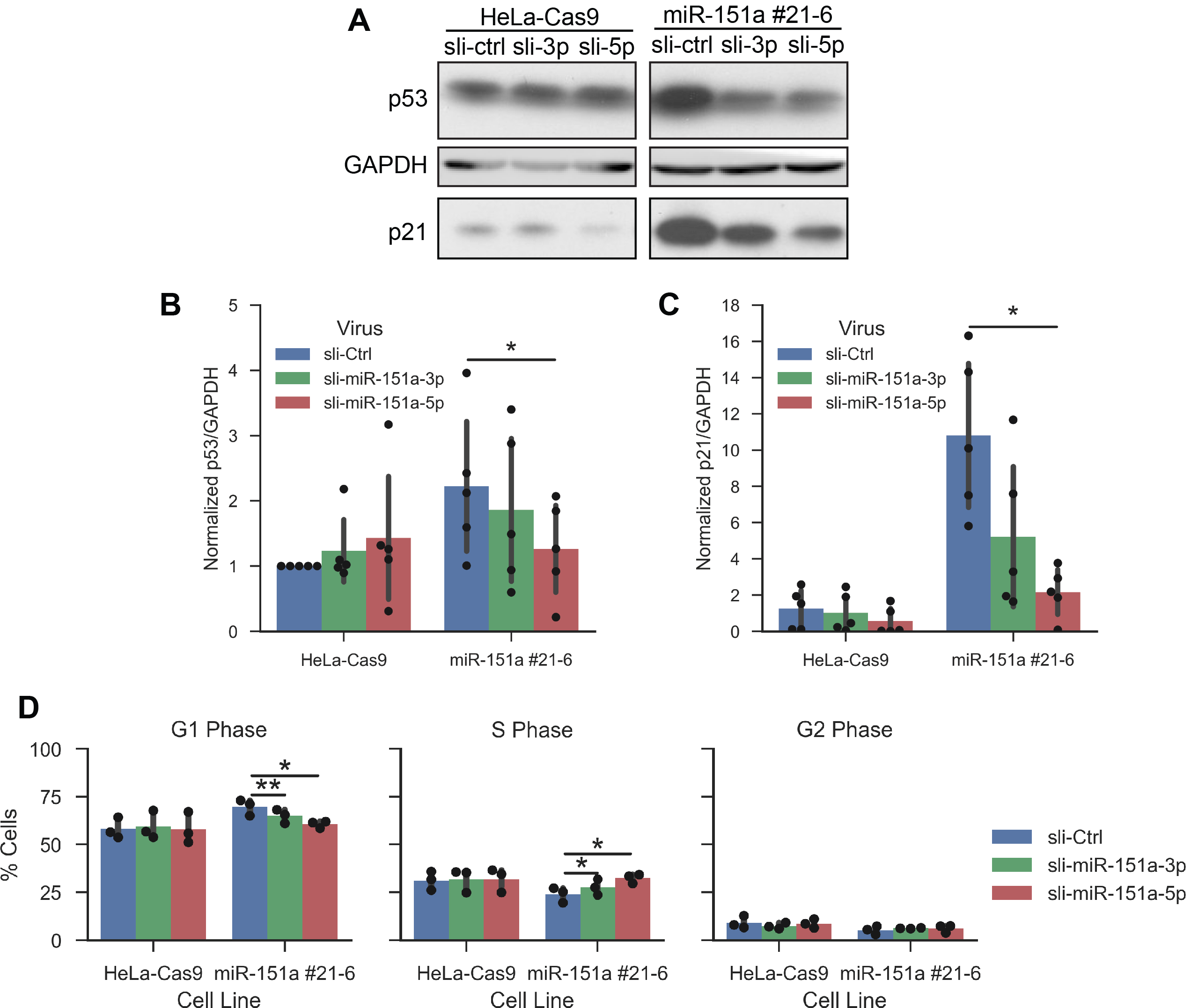
Revert molecular and cellular phenotypes in mutant cells after re-expression of miR-151a. **A**, Representative western blot of p21 and p53 levels in HeLa-Cas9 parental and miR-151a #21-6 mutant cells transduced with sli-Ctrl, sli-miR-151a-3p, or sli-miR-151a-5p. GAPDH was used as a loading control. **B**, Quantification of p53 over GAPDH levels and normalized to the ratio in HeLa-Cas9 transduced with sli-Ctrl. Data points represent five independent experiments. **C**, Quantification of p21 over GAPDH and normalized to the mean value in HeLa-Cas9 transduced with sli-Ctrl. Data points represent five independent experiments. **D**, HeLa-Cas9 and miR-151a #21-6 cells transduced with sli-Ctrl, sli-miR-151a-5p, or sli-miR-151a-3p were incubated in BrdU-containing medium, then co-stained with anti-BrdU and PI and analyzed by flow cytometry to determine the cell cycle distribution. Data points represent three independent experiments. For **B**-**D**, statistics were calculated using two-tailed paired *t*-test (* *p* < 0.05; ** *p* < 0.01). Error bars represent 95% confidence interval.

To determine if the increase in G1 phase cells seen after miR-151a knockout can be reversed, we examined the cell cycle in the re-expressing cells. We found in transduced miR-151a #21-6 cells, re-expression of miR-151a-5p or -3p was able to decrease the percentage of cells in G1 and to increase cells in the S phase (Fig. 5D). The decrease in G1 was greater after miR-151a-5p re-expression than after miR-151a-3p re-expression. No significant changes were observed in miR-151a #14 cells transduced with either sli-miR-151a-5p or -3p (Supplemental Fig. S5D). Although the results from miR-151a mutant clone #14 with partial loss-of-function alleles were somewhat complexed, the results from miR-151a complete knockout clone #21-6 supported the notion that miR-151a, especially the 5p miRNA, regulated p53/p21 and the cell cycle.

In total, we found loss of miR-151a resulted in a decrease in cellular fitness and an increased arrest in the G1 phase. miR-151a knockdown increased the p53 and p21 protein levels. Re-expression of either miR-151a-5p or miR-151a-3p suppressed p53 and p21; however, miR-151a-5p apparently did not target p53 mRNA directly, while 3p could directly target the 3’-UTR of p53. Re-expression of miR-151a-5p was able to reverse the G1 arrest; re-expression of 3p was able to reverse as well, but to a less extent.

### miR-151a and p21 in Patient Samples

To investigate if the miR-151a-p53-p21 regulation is of clinical relevance, we examined the RNA expression level of miR-151a, p53, and p21 in patient samples. Upon examining the data of the Cancer Genome Atlas (TCGA) cervical squamous cell carcinoma and endocervical adenocarcinoma (CESC) project, no significant negative correlation between miR-151a and TP53 was found (Supplemental Fig. S6). However, we found a significant negative correlation between miR-151a-5p expression and p21/CDKN1A expression (Fig. 6A). Also, there appeared to be a negative correlation between miR-151a-3p and CDKN1A expression; however, it did not reach significance (p-value: 0.065; Fig. 6B).

**Figure 6.**
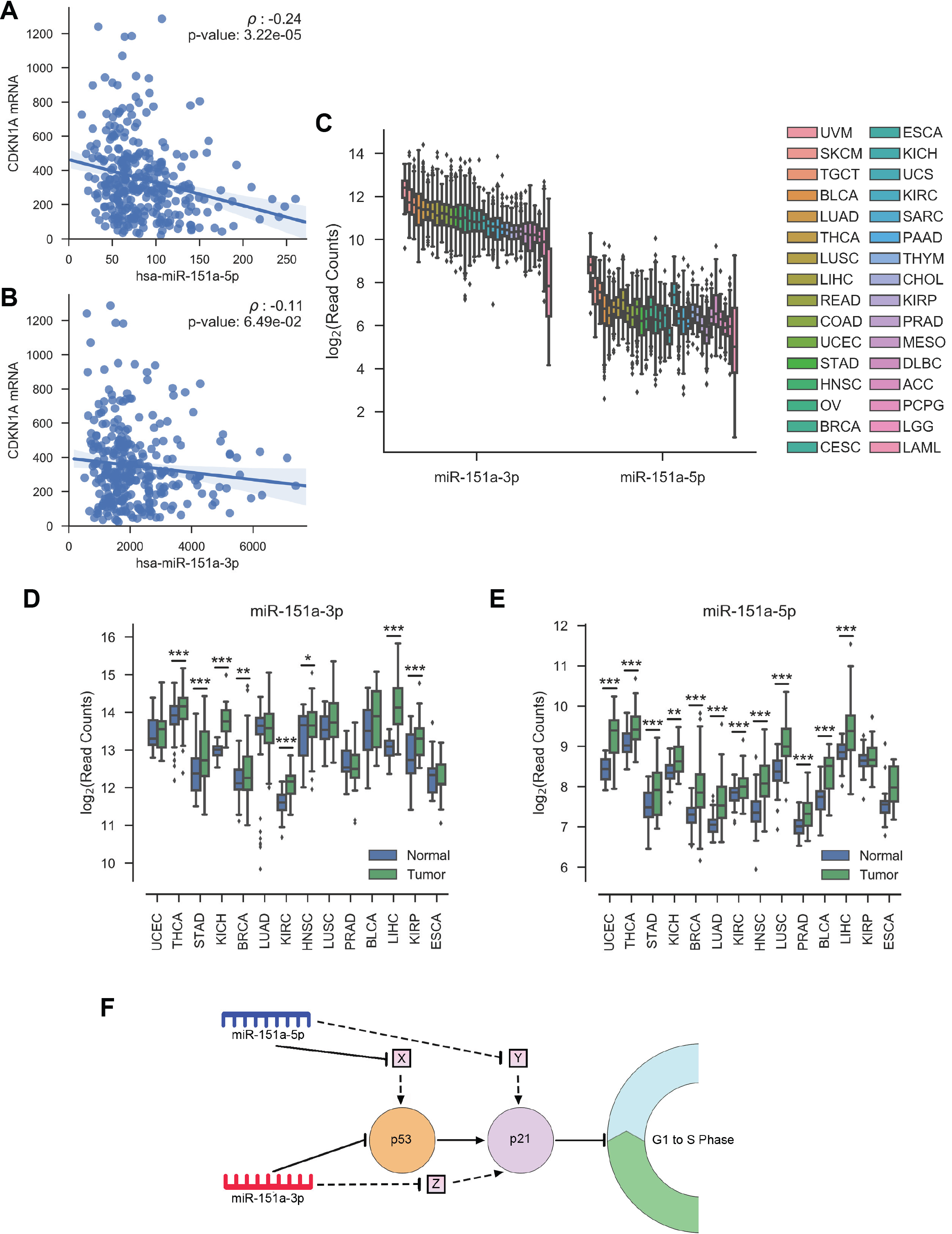
Expression of miR-151a in patient samples and its correlation with CDKN1A/p21 in tumors. **A, B**, The Pearson’s correlation coefficient between **A**, miR-151a-5p and CDKN1A, or **B**, miR-151a-3p and CDKN1A in CESC patient samples was determined. The line indicates the linear regression and the shaded area represents the 95% CI. **C**, The box plot shows miR-151a expression in 32 different tumor types based on RNA-Seq data in TCGA. **D, E**, The expression of **D**, miR-151a-3p or **E**, miR-151a-5p in paired tumor and normal samples was examined. The significance of the difference in expression was calculated using DESeq2 (* *p* < 0.05; ** *p* < 0.01; *** *p* < 0.001). For **C-E**, the box shows the quartiles, the lines show the range, and dots represent outliers. **F**, A proposed model of miR-151a regulation of p53, p21, and the cell cycle. A connecting line ends with a bar indicates suppression, while a line ends with an arrowhead indicates activation. Experimentally supported regulation is depicted with a solid line, while proposed regulation is depicted with a broken line. X, Y, and Z are hypothetical intermediate regulators.

We then broadened our scope and examined the expression of miR-151a across many cancers. miR-151a is detected in all 32 TCGA cancer types examined, with higher miR-151a-3p than miR-151a-5p expression in all samples (Fig. 6C). When we examined the 14 cancers for which TCGA has a significant number of paired tumor and normal tissue samples, we found the expression of miR-151a-3p was significantly increased in 8 cancers: thyroid (THCA), stomach (STAD), kidney chromophobe (KICH), breast (BRCA), kidney renal clear cell (KIRC), head/neck (HNSC), liver (LIHC), and kidney renal papillary (KIRP) cell tumors compared to normal tissues (Figure 6D), while the expression of miR-151a-5p was significantly increased in 12 cancers, including uterine corpus endometrial (UCEC), lung adenocarcinoma and squamous cell carcinoma (LUAD and LUSC), prostate (PRAD), bladder (BLCA), and 7 of the aforementioned tumor types (Fig. 6E). In total, 13 of the 14 cancers examined show increased expression of miR-151a-5p, miR-151a-3p, or both in tumor samples (Table 1). None of 14 cancer types showed a significant decrease in miR-151a-3p or -5p expression. This indicated miR-151a up-regulation may be a common event in tumor formation.

**Table 1.**
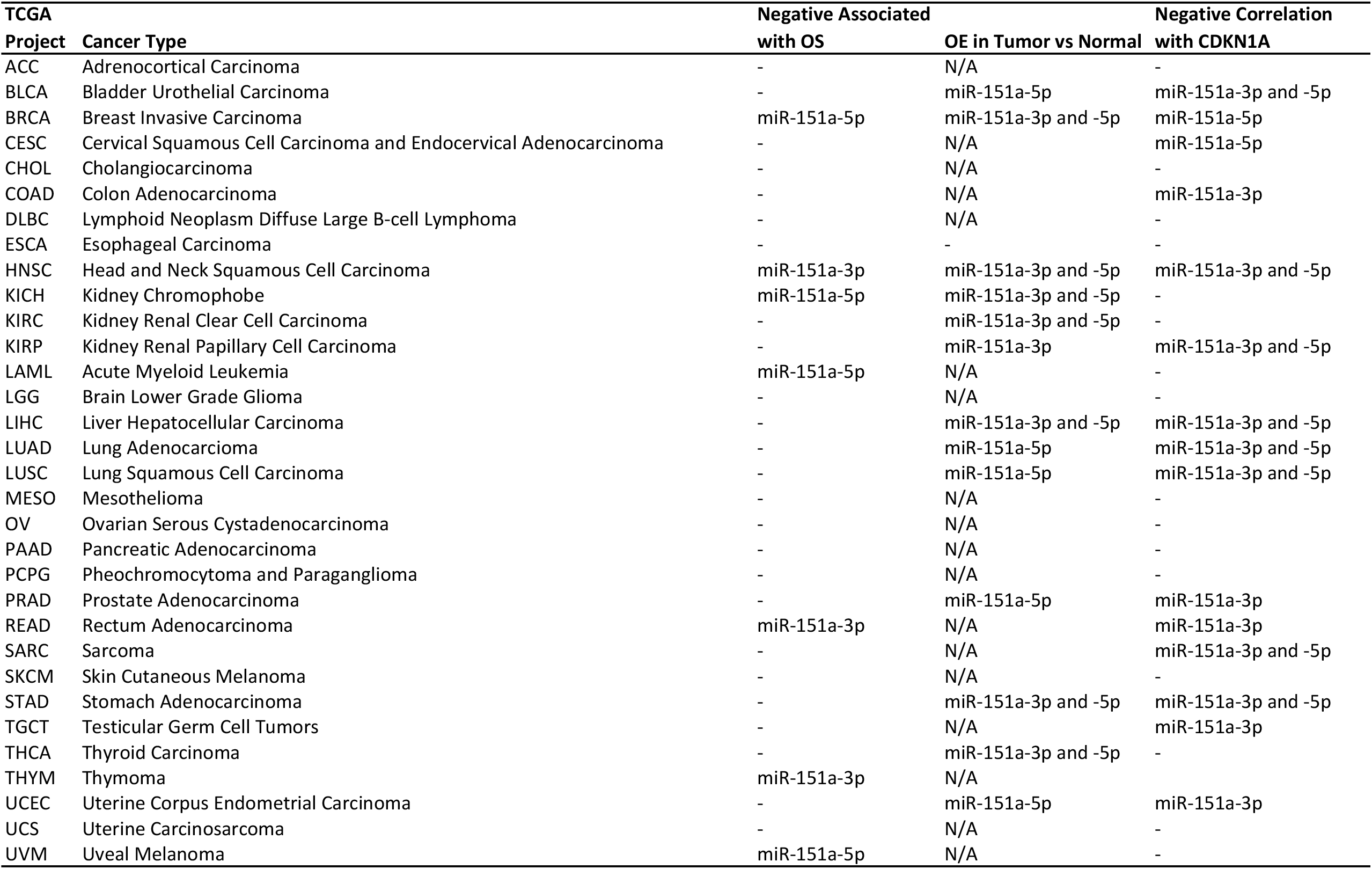
TCGA data summary

We also looked at the association of miR-151a expression with overall survival (OS). When we examined the 25% of patients with the highest versus the 25% with lowest miR-151a expression, we found high miR-151a-5p expression is associated with decrease OS in 4 cancers: kidney chromophobe (KICH), uveal melanoma (UVM), breast carcinoma (BRCA), and acute myeloid leukemia (LAML) (Table 1). High miR-151a-3p expression is linked to decreased OS in 3 cancers: sarcomas (SARC), head and neck (HNSC), and thymoma (THYM) (Table 1). In no case that a high expression of miR-151a-5p or -3p was found to associate with increased OS Thus, in 7 of the 32 cancers examined, patients with a high expression of miR-151a have decreased OS compared to those with a relatively low miR-151a expression.

As we found a negative correlation between miR-151a and p21/CDKN1A expression in cervical cancer, we wanted to examine this relationship in other cancer patient samples in TCGA. We found a significant inverse correlation between miR-151a (−5p, -3p, or both) and CDKN1A in 14 more of the 32 cancer types (Table 1). Only a single cancer type, thymoma (THYM), was an exception which had a positive correlation between miR-151a expression and CDKN1A expression. The negative correlation seen in 15 cancers suggested miR-151a’s suppression of p21 expression and dysregulation of the cell cycle may be clinically relevant.

## DISCUSSION

In this study, we examined miR-151a in cellular fitness and in cancer correlation. We used CRISPR-Cas9 to mutate the miR-151a gene in HeLa cells and found loss of miR-151a expression resulted in a decrease in cell growth and an increase of cell population at the G1-phase of the cell cycle. By analyzing transcriptome changes, we found genes involved in the cell cycle, p53, and TGF-β pathways were disproportionally up-regulated in miR-151a knockout cells. We also found miR-151a knockout resulted in an increase in p53 and p21 protein expression. miR-151a-3p directly, while -5p indirectly suppressed p53-3’UTR reporters. miR-151a-5p re-expression was able to decrease the p53 and p21 protein levels as well as decrease the percentage of G1-phase cells, while miR-151a-3p re-expression only modestly decreased the p53 or p21 protein level. From the TCGA database, we found nearly half of the thirty-two cancers examined, including cervical cancer, had a negative correlation between miR-151a and p21/CDKN1A expression. Together, these results suggested that suppression of p21 through miR-151a overexpression may be important for tumorigenesis or progression in a broad range of cancers.

The reporter and miRNA re-expression experiments indicated direct targeting of p53 by miR-151a-3p is not the primary mechanism by which miR-151a regulates p53 and p21 expression. It has been reported that p53 is a miR-151a target in gastric cancer cells where overexpression of pre-miR-151a, which can produce both mature miR-151a-5p and -3p, decreases luciferase expression from a construct containing the 3’-UTR of TP53 ^7^. They attributed this regulation to miR-151a-5p due to the presence of a RNAhybrid-predicted binding site in the 3’-UTR of TP53 and the greater suppressive effects of anti-miR-151a-5p on cell migration and invasion ^7^. However, our study and others ^37^ indicate that it is miR-151a-3p that directly suppresses p53 3’UTR, not miR-151a-5p. It has been postulated that a number of microRNAs can target TP53 gene directly or indirectly through other regulators in the p53 network ^38^. Nonetheless, all these studies support the notion that miR-151a indeed regulates p53 expression, although it is still not clear how miR-151a-5p regulates the expression of p53 and p21. A confirmed target of miR-151a-5p is *ARHGDIA* ^4, 5^, which suppresses Rho activation by preventing the dissociation of GDP. Since Rho can suppress p21^39^, miR-151a-5p could suppress p21 through suppression of ARHGDIA which leads to increased Rho signaling. Our RNA-Seq data did not show significant increases of *ARHGDIA* mRNA level in miR-151a mutant cells, but we could not rule out the possibility that miR-151a-5p suppresses *ARHGDIA* primarily at the translational level without drastically altering its mRNA stability.

FAK/PTK2 is encoded by the host gene of miR-151a, which can also negatively regulate p53. By binding to p53, FAK inhibits p53’s ability to activate transcription and induces p53 degradation ^40, 41^. Interestingly, p53 can transcriptionally regulate miR-151a and FAK expression. There are two p53-responsive elements located up-stream of miR-151a, and miR-151a expression is higher in p53^+/+^ than p53^-/-^ HCT116 cells ^42^, indicating p53 may induce miR-151a expression through a promoter near the p53-responsive elements. p53 can also regulate the processing of pri-miR-151a, as p53 has been shown to associate with Drosha and DDX5 to enhance miRNA processing ^43^. Furthermore, p53 is able to repress the transcription of the FAK locus ^44^, and knockdown of p53 by siRNA has been shown to increase both levels of FAK and miR-151a in HepG2 cells ^4^. Thus, there appears to be a complicated network of interactions: both FAK and miR-151a suppress p53, p53 suppresses FAK, and p53 promotes or suppresses miR-151a that may be cellular context dependent.

Along with p53 and cell cycle pathway genes, we also saw an enrichment of TGF-β pathway genes among those up-regulated in miR-151a knockout mutants. There are three members in the TGF-β family of growth factors in humans: TGF-β1, TGF-β2, and TGF-β3. Of these three, both TGFB3, which encodes TGF-β3, and TGFB2, which encodes TGF-β2, were significantly up-regulated in our mutant cells. TGFB3 is a predicted miR-151a-5p target, but there is no evidence TGFB2 is a direct target of miR-151a. We did not detect an increase of the TGFB1 mRNA, which encodes TGF-β1, in our mutants. Quantification based on our mRNA-Seq data suggested the mRNA level of TGFB3 and TGFB2 were approximately one thirtieth to one ninth of that of TGFB1, so it is unclear if up-regulation of B3 and B2 was sufficient to affect TGF-β signaling. Several TGF-β-related genes are also miR-151a-3p predicted targets, including the TGF-β receptor genes TGFBR1 and TGFBR2, though the mRNA levels of these genes did not significantly change in our mutant cells. It would be of interest to further investigate the potential changes to TGF-β signaling upon knocking out of miR-151a, since this pathway has been shown to up-regulate p21 through SMAD-mediated transcriptional regulation of *CDKN1A* ^45^.

We propose a model to summarize the key observations (Fig. 6F). When miR-151a is up-regulated in tumor cells, miR-151a-3p directly suppresses p53, while miR-151a-5p indirectly suppresses p53. The down-regulation of p53 prevents the transcriptional activation of p21 and p21-mediated G1-arrest. miR-151a-3p and miR-151a-5p may also suppress p21 in a p53-independent manner, as knockdown of p53 in miR-151a mutant cells did not fully return the p21 protein level back down to wild type levels. It is imperative to identify those uncharacterized miR-151a targets involved in these regulations. We also propose that miR-151a’s regulation of the cell cycle through p53 and p21 may be of clinical relevance, since in many cancers miR-151a is overexpressed, the p21 level is negatively correlated with the miR-151a level, and a high miR-151a level is associated with poor prognosis. It is worth pointing out that both 5p and 3p of miR-151a may work cooperatively to regulate the p21 pathway. Since the ratio of the miRNA-5p and the miRNA-3p generated from a microRNA gene can be important for cancer ^46^, further investigation of the combinatorial effect of miR-151a-5p and -3p on tumorigenesis shall be of interest.

## ACKNOWLEDGEMENTS

We thank City of Hope colleagues Jiing-Kuan Yee for plasmids, Xiwei Wu, Jinhui Wang, and Yate-Ching Yuan for help with RNA-seq and bioinformatics, and Jeremy Stark and Lucy Brown for flow cytometry and cell cycle analysis. We also thank JSK’s dissertation committee for valuable suggestions and comments and QiWen Li and Willow Huang of the summer Roberts Academy for technical and graphic assistance. This work was supported by Beckman Research Institute grants to RJL, and by NIH grant P30-CA033572 for shared research core facilities at City of Hope.

## AUTHORS’ CONTRIBUTIONS

JSK and RJL conceived the project and designed experiments; JSK performed experiments; JSK and RJL analyzed the data and wrote the paper.

