## Supplemental material for "Pan-Cancer Biomarker miR-151a Regulates p21 Partially Through p53"

Supplemental Table S1: Oligo Sequences

| Primer Name | Oligo Sequence |
|---|---|
| sg-151a-F | CACCGGGGATGGTCATACTCACCT |
| sg-151a-R | AAACAGGTGAGTATGACCATCCCC |
| miR-151a-FA-F | TGTAAAACGACGGCCAGTATTCAGTGCCTGGGTGACTC |
| miR-151a-FA-R | AGCTGAGCCTGGTGCTAGTC |
| M13(-21)-FAM | FAM-TGTAAAACGACGGCCAGT |
| sli-miR1515p-F | gatcaCGAGGAGCTCACAGTCTAGTaAGACTGTGAGCTCCTCGcTTTTT |
| sli-miR1515p-R | tcgaAAAAAgCGAGGAGCTCACAGTCTtACTAGACTGTGAGCTCCTCGt |
| sli-miR1513p-F | gatcaTAGACTGAAGCTCCTTGAGGaCAAGGAGCTTCAGTCTAcTTTTT |
| sli-miR1513p-R | tcgaAAAAAgTAGACTGAAGCTCCTTGtCCTCAAGGAGCTTCAGTCTAt |
| sli-Ctrl-F | gatcaGCGTTCTACACTCGACGTACTtGTCGAGTGTAGAACGCcTTTTT |
| sli-Ctrl-R | tcgaAAAAAgGCGTTCTACACTCGACAAGTACGTCGAGTGTAGAACGCt |
| TP53-UTR-F | AATTCTAGGCGATCGCTCGACCACCTGAAGTCCAAAAAGG |
| TP53-UTR-R | GCGGCCAGCGGCCGCGTCGAAAATGCAGATGTGCTTGCAG |
| TGFB3-UTR-F | AATTCTAGGCGATCGCTCGATCCAACATGGTGGTGAAGTC |
| TGFB3-UTR-R | GCGGCCAGCGGCCGCGTCGAccaggatgccccaaaaata |
| TP53-3p-SDM-F | agttttttatggcgggaggtactgtgacccttttggacttcagg |
| TP53-3p-SDM-R | cctgaagtccaaaaagggtcacagtacctcccgccataaaaaact |
| TP53-5p-SDM-F | acaggtggatcgcctgacgacaggagtttgagaccag |
| TP53-5p-SDM-R | ctggtctcaaactcctgtcgtcaggcgatccacctgt |
| TP53-5p2-SDM-F | cgcctgagccctccagtttgagaccagcctgggca |
| TP53-5p2-SDM-R | tgcccaggctggtctcaaactggagggctcaggcg |
| TGFB3-5p-SDM-F | gtgtgtttcccgtccagcgggcagtcaggcagt |
| TGFB3-5p-SDM-R | actgcctgactgcccgctggacgggaaacacac |
| sh-p53-F | AAAAGACTCCAGTGGTAATCTACTTCAAGAGAGTAGATTACCACTGGAGTC |
| sh-p53-R | AAAAGACTCCAGTGGTAATCTACTCTCTTGAAGTAGATTACCACTGGAGTC |

Red indicates primer tails/adaptor sequences

**Table S2.** miR-151a predicted targets up-regulated and related to cell cycle, p53, or TGFβ

| Gene Symbol | GO term(s) | Category | Targeted by |
|---|---|---|---|
| AZI2 | mitotic cell cycle | Cell Cycle | miR-151a-3p |
| BCL2 | negative regulation of G1/S transition of mitotic cell | Cell Cycle | miR-151a-5p |
| CCND2 | cell cycle; positive regulation of G1/S transition of mitotic cell cycle | Cell Cycle | miR-151a-3p |
| CCPG1 | cell cycle | Cell Cycle | miR-151a-3p |
| DUSP1 | negative regulation of meiotic cell cycle; mitotic cell cycle arrest; regulation of mitotic cell cycle spindle assembly checkpoint | Cell Cycle | miR-151a-3p and -5p |
| EGF | positive regulation of mitotic nuclear division | Cell Cycle | miR-151a-3p |
| GAS7 | cell cycle arrest | Cell Cycle | miR-151a-3p |
| GATA3 | negative regulation of cell cycle | Cell Cycle | miR-151a-5p |
| INSR | positive regulation of meiotic cell cycle; positive regulation of mitotic nuclear division | Cell Cycle | miR-151a-3p and -5p |
| KIF3B | mitotic spindle organization; mitotic centrosome separation; mitotic spindle assembly | Cell Cycle | miR-151a-3p |
| PAK3 | regulation of mitotic cell cycle | Cell Cycle | miR-151a-3p |
| PDGFRB | positive regulation of mitotic nuclear division | Cell Cycle | miR-151a-5p |
| PELO | cell cycle | Cell Cycle | miR-151a-3p |
| PPP2R2C | mitotic cell cycle | Cell Cycle | miR-151a-5p |
| TP53 | positive regulation of cell cycle arrest; regulation of cell cycle G2/M phase transition; positive regulation of cell cycle arrest; mitotic G1 DNA damage checkpoint; DNA damage response, signal transduction by p53 class mediator; DNA damage response, signal transduction by p53 class mediator resulting in cell cycle arrest; DNA damage response, signal transduction by p53 class mediator resulting in transcription of p21 class mediator; intrinsic apoptotic signaling pathway in response to DNA damage by p53 class mediator; signal transduction by p53 class mediator; intrinsic apoptotic signaling pathway by p53 class mediator; regulation of signal transduction by p53 class mediator | Cell Cycle; p53 | miR-151a-3p |
| SOX9 | regulation of cell cycle process; cellular response to transforming growth factor beta stimulus | Cell Cycle; TGF-β | miR-151a-3p |
| THBS1 | cell cycle arrest; positive regulation of transforming growth factor beta receptor signaling pathway; positive regulation of transforming growth factor beta1 [illegible] | Cell Cycle; TGF-β | miR-151a-3p |
| MAPK11 | regulation of signal transduction by p53 class mediator | p53 | miR-151a-5p |
| COL1A1 | cellular response to transforming growth factor beta stimulus | TGF-β | miR-151a-3p |
| LDLRAD4 | negative regulation of transforming growth factor beta receptor signaling pathway | TGF-β | miR-151a-3p and -5p |
| ONECUT2 | negative regulation of transforming growth factor beta receptor signaling pathway | TGF-β | miR-151a-3p |

| | | | |
|---|---|---|---|
| SMAD9 | transforming growth factor beta receptor signaling pathway | TGF-β | miR-151a-3p |
| TGFB3 | transforming growth factor beta receptor signaling pathway; negative regulation of transforming growth factor beta receptor signaling pathway | TGF-β | miR-151a-5p |

**Supplementary Table S3.** Significantly increased in p53 consensus targets in miR-151a mutants

| Gene Name | $log_2$(Fold Change) | Corrected p-value | Known Function |
|---|---|---|---|
| CDKN1A | 1.73 | 1.27E-06 | Cell Cycle |
| ENC1 | 2.25 | 6.82E-06 | Proliferation |
| POLH | 0.79 | 1.16E-05 | DNA Damage Response |
| HHAT | 4.42 | 9.23E-05 | Proliferation |
| PHLDA3 | 0.68 | 0.00036 | AKT Regulation |
| RPS27L | 0.63 | 0.00058 | DNA Damage Response; Cell Cycle |
| DDB2 | 0.81 | 0.00083 | DNA Damage Response |
| PLK2 | 1.17 | 0.00095 | Proliferation |
| BBC3 | 1.21 | 0.00228 | Apoptosis |
| TIGAR | 0.70 | 0.00257 | Apoptosis Inhibition |
| GDF15 | 4.35 | 0.00287 | TGF-β Signaling |
| FBXO22 | 0.47 | 0.00407 | Ubiquitination |
| AEN | 0.51 | 0.00599 | Apoptosis |
| SAC3D1 | 0.54 | 0.00715 | Cell Cycle |
| BAX | 0.53 | 0.00876 | Apoptosis |
| FAM13C | 1.24 | 0.04425 | - |

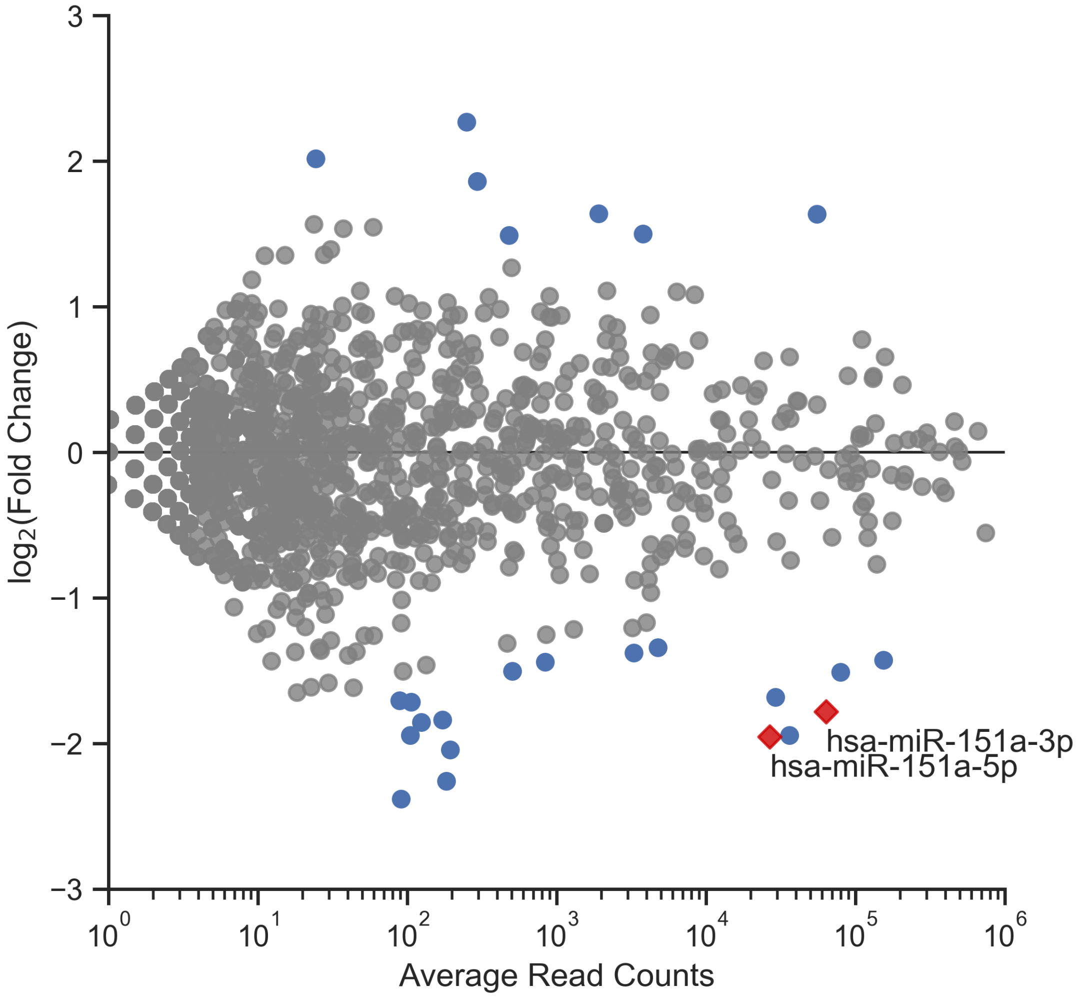


**Supplementary Figure S1**. miRNA-sequencing of miR-151a #14 and HeLa-Cas9. Small RNA from HeLa-Cas9 parental and miR-151a #14 mutant cells was collected once and sequenced. Blue indicates miRNAs with raw *p* < 0.05. Red diamonds indicate miR-151a-3p and -5p. p-value calculated using DESeq2.

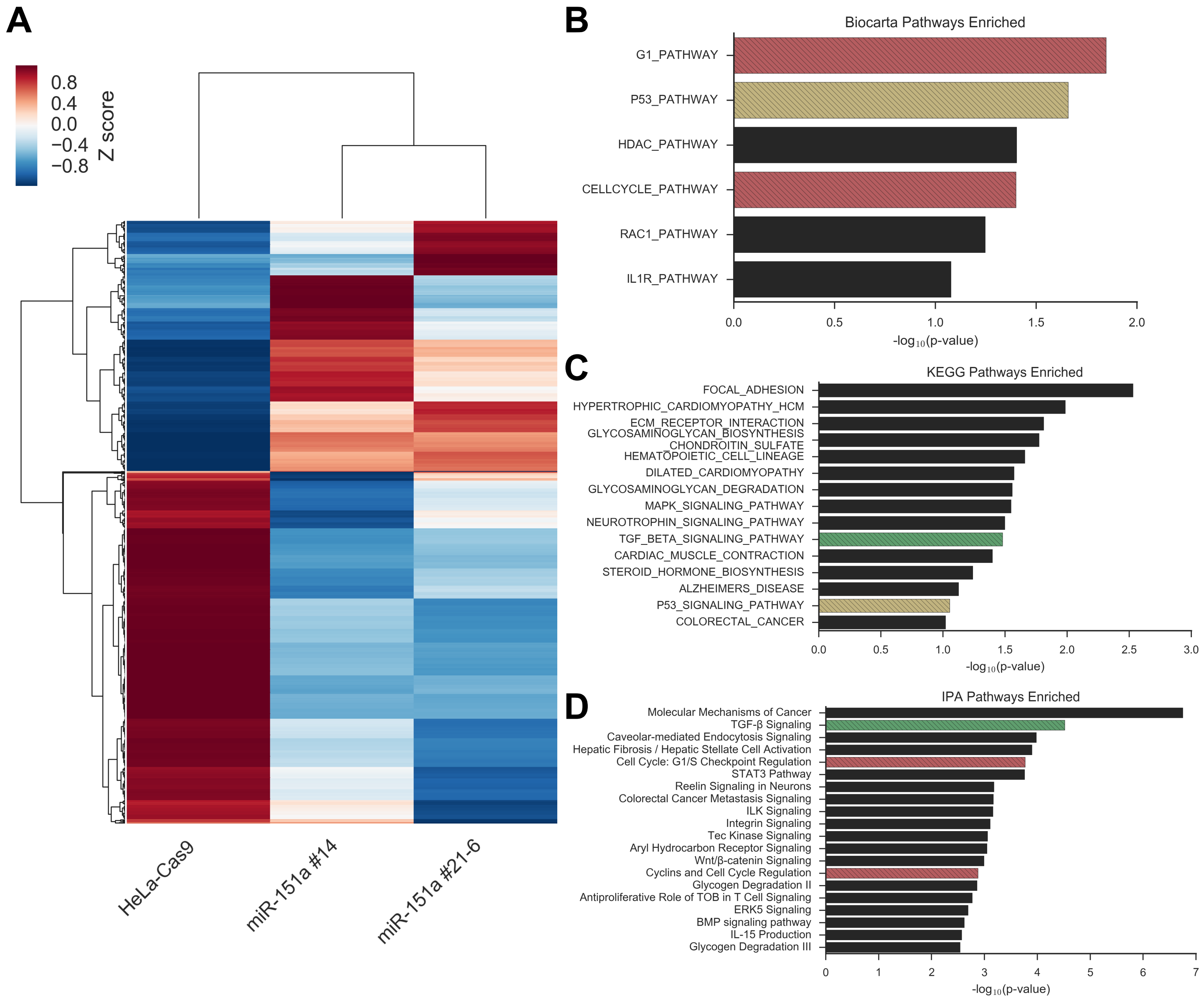


**Supplementary Figure S2**. mRNA-sequencing analysis. **A,** The mRNA expression profiles of HeLa-Cas9 and miR-151a mutant cells was determined by mRNA-sequencing. The heat map shows the z-score normalized mRNA-sequencing reads for EdgeR significantly differentially expressed (FDR < 0.05) mRNAs. The read counts are the average of two independent replicates. **B-D,** The **B,** Biocarta **C,** KEGG and **D,** Ingenuity pathways enriched among up-regulated mRNAs was determined using **B,C,** Gene Set Enrichment Analysis or **D,** Ingenuity Pathway Analysis. For **B-D**, cell cycle related pathways are shown in red, p53 related pathways are in yellow and TGFβ related pathways are in green.

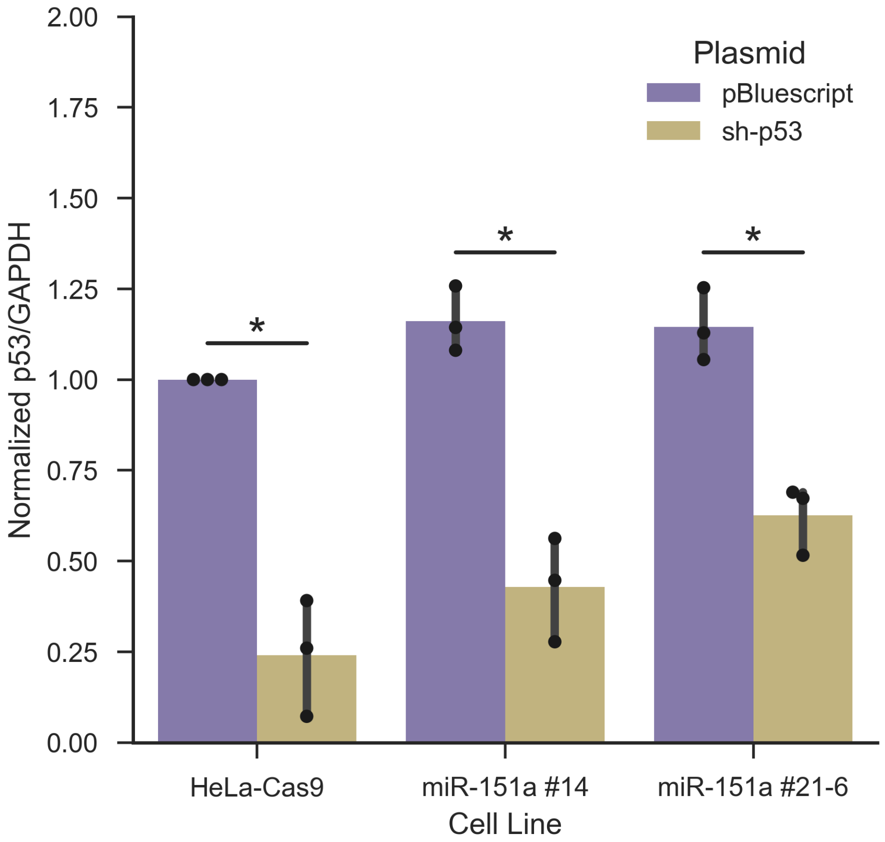


**Supplementary Figure S3**. p53 expression after sh-p53 transfection. The p53 expression level in HeLa-Cas9 parental and miR-151a mutant cells after transfection with sh-p53 or empty vector was determined by western blot. Data points indicate three biological replicates. Significance was determined using the two-tailed paired t-test (* $p < 0.05$). Error bars represent 95% confidence interval.

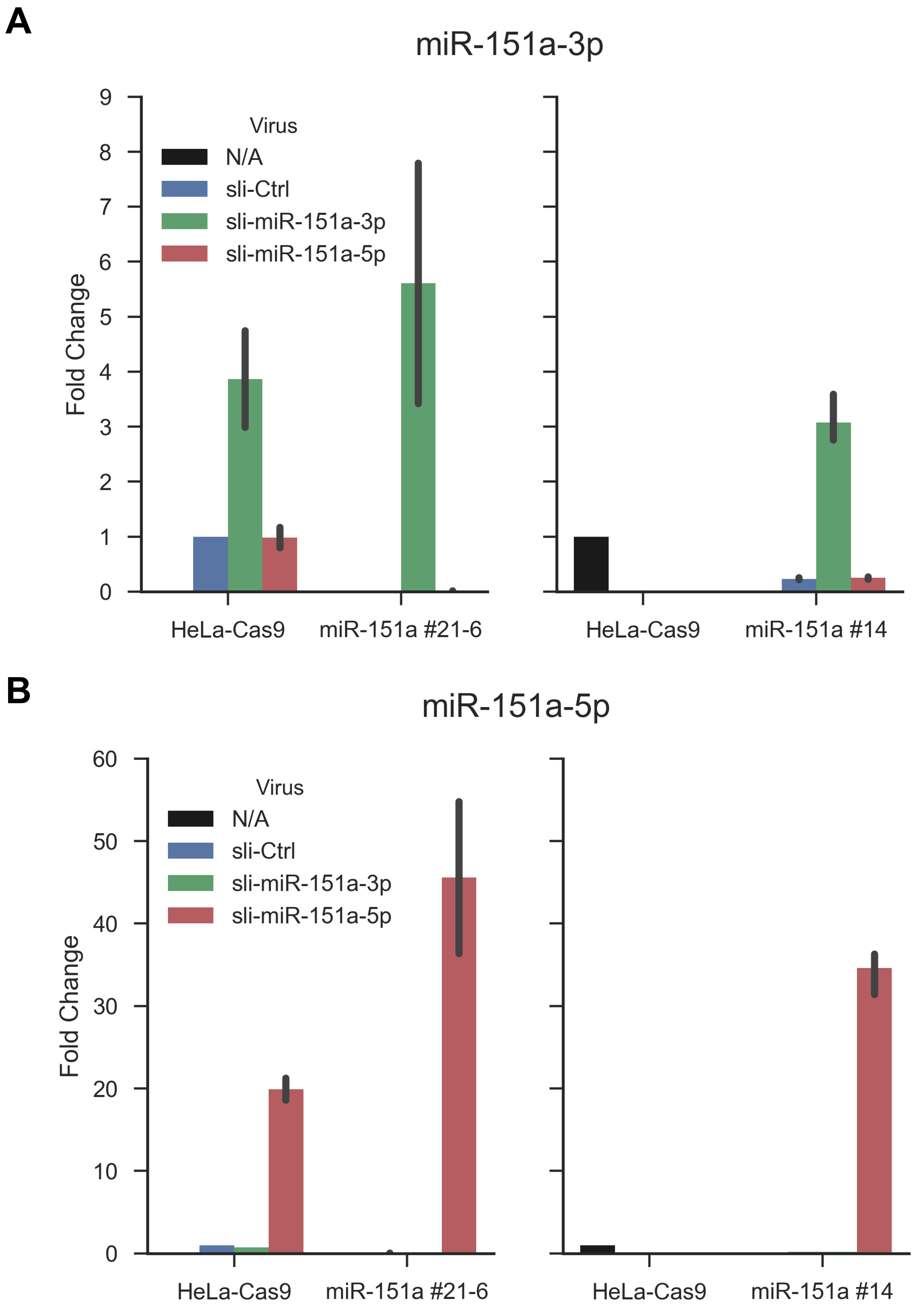


**Supplementary Figure S4**. miR-151a expression after sli-miR transduction. The **A,** miR-151a-3p and **B,** miR-151a-5p expression level was measured by qRT-PCR after transduction of HeLa-Cas9 parental or miR-151a mutant cells with a slicer scramble control (sli-ctrl) or slicer miR-151a-3p or -5p overexpression constructs. The HeLa-Cas9 and colony miR-151a #21-6 values were normalized to HeLa-Cas9 sli-ctrl cells. Colony miR-151a #14 values were normalized to HeLa-Cas9 untransduced cells.

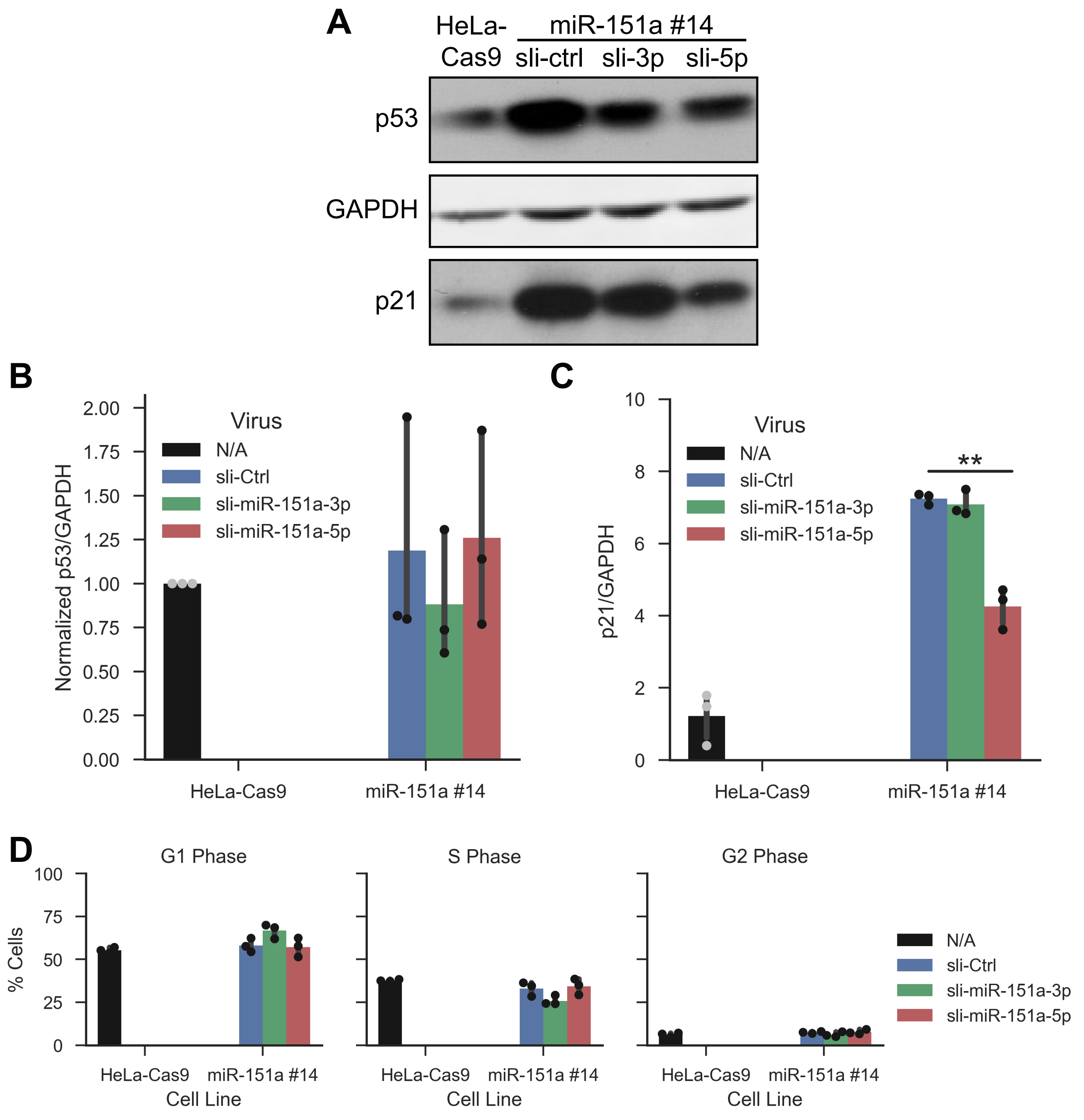


**Supplemental Figure S5.** miR-151a re-expression in colony miR-151a #14. **A,** Representitive western blot of p21 and p53 for HeLa-Cas9 parental and miR-151a #14 cells transduced with sli-miR-151a-5p, -3p or sli-Ctrl. **B,** Quantification of p53 levels in transduced miR-151a #14 cells normalized to GAPDH and HeLa-Ca9 parental expression levels. **C,** Quantification of p21 levels in transduced miR-151a #14 cells normalized to GAPDH and the mean HeLa-Ca9 parental expression level. **D,** Cells were incubate with BrdU-containing medium, co-stained with anti-BrdU and PI and analyzed by flow cytometry to determine the cell cycle distribution. For **B**-**D**, statistics were calculated using two-tailed paired t-test (* $p < 0.05$; ** $p < 0.01$). Error bars represent 95% confidence interval. Data points represent three independent experiments.

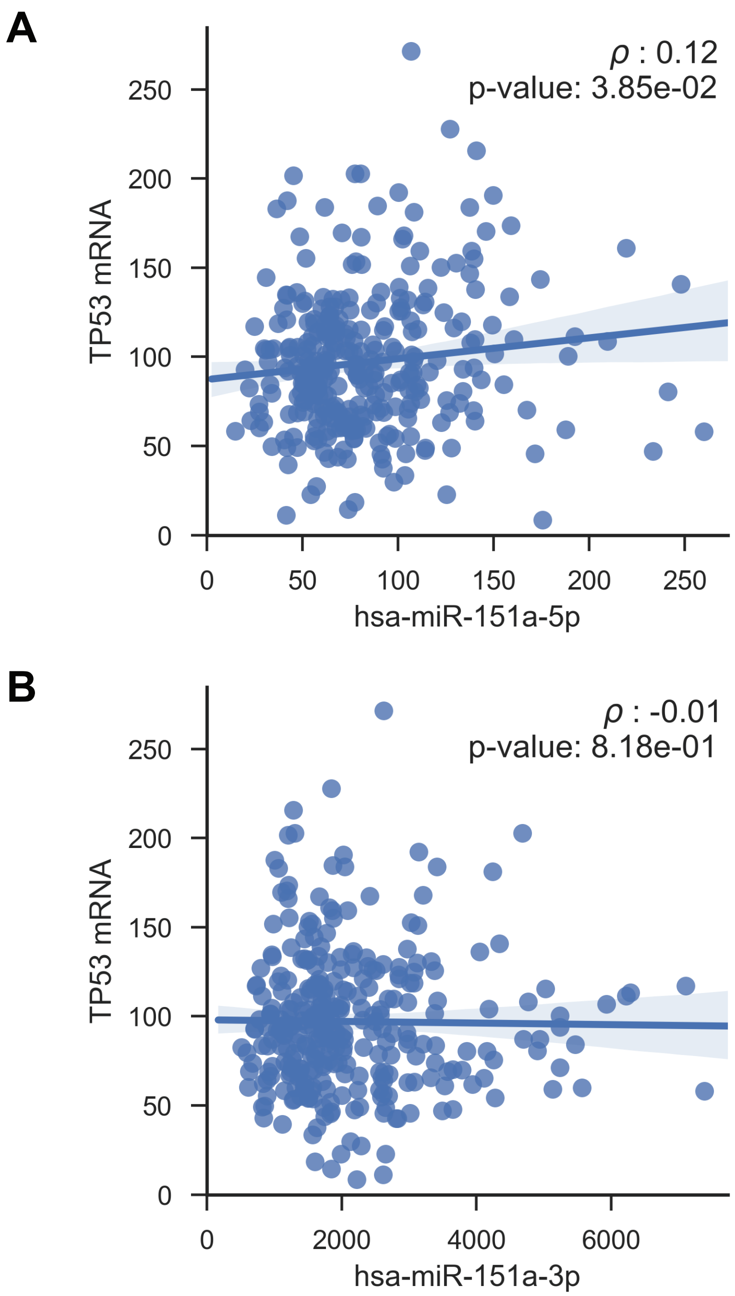


**Supplementary Figure S6**. Correlation of miR-151a and TP53 expression in CESC patient samples. The Pearson's correlation coefficient between **A,** miR-151a-5p or **B,** miR-151a-3p and TP53 expression was determined for CESC patient samples. The line represents the linear regression and the shaded area represents the 95% CI for the linear regression.
